# An *in-silico* approach to study the molecular interaction of the PINK1 kinase domain with Parkin Ubl domain

**DOI:** 10.1101/2023.05.02.539044

**Authors:** Sima Biswas, Sreekanya Roy, Angshuman Bagchi

**Affiliations:** Department of Biochemistry and Biophysics, University of Kalyani, Kalyani, Nadia-741235, West Bengal, India

**Author notes:** Corresponding author Angshuman Bagchi. E-mail: Sima Biswas.

**Keywords:** PINK1, Parkin, Parkinson’s Disease, Mutations, Docking, MD simulation

## Abstract

Parkinson’s Disease (PD) is the second most common neurodegenerative disorder which creates devastating effects on the neurons of the mid brain. PARK2 and PARK6 genes encode the E3 ubiquitin ligase Parkin and serine/threonine protein kinase PINK1 (PTEN-induced kinase 1), respectively. Mutations in these proteins are responsible for majority of the early onset of PD. PINK1 protein along with Parkin are known to participate in the mitochondrial mitophagy pathway which selectively removes damaged mitochondria. Parkin with the help of its intra-domain interactions goes to an autoinhibited inactive state. Parkin ubiquitin activity is suppressed in this condition. Phosphorylation of the Serine65 residue, present in the Ubl domain of Parkin, is required for its ubiquitin ligase activity. PINK1 protein phosphorylates the Serine65 residue after its self-phosphorylation. After activation, Parkin protein ubiquitinates other damaged mitochondrial proteins for their degradation. Therefore, PINK1-Parkin interaction is important for the proper maintenance of the mitochondrial and neuronal fidelity. In this present work, we are trying to find out how the mutations in the PINK1 kinase domain would affect the modes of its interaction with the Ubl domain of Parkin. We used the three dimensional coordinates of the PINK1 and Ubl domain of Parkin from our previous work and selected the mutations P296L and G309D which have the abilities to hamper the said interactions between PINK1 and the Ubl domain of Parkin. We used molecular docking followed by molecular dynamics simulations to study the nature and structural dynamics of the binding interactions. The results from our study could provide an insight into the PINK1 mediated Parkin activation and plausible biochemical mechanism behind the onset of PD.

## INTRODUCTION

Autosomal Recessive Juvenile Parkinson’s Disease (ARJP) is caused due to the mutations in the PINK1 and Parkin proteins (Kitada et al., 1998) (Cookson, 2012). Mutations in these two proteins are responsible for the Familial form of Parkinson’s Disease (PD). PINK1, encoded by PARK6 gene, a PTEN-induced putative kinase 1 and Parkin, encoded by PARK2 gene, an E3 ubiquitin ligase, work together along with other proteins in mitochondrial quality control pathway (Quinn et al., 2020) (Agarwal & Muqit, 2022). These two proteins are very important for the damaged mitochondrial ubiquitination and eventually their degradations (Deas et al., 2011) (Ge, Preston et al., 2020). In normal cellular condition, PINK1 protein is continuously processed and degraded by proteasomal degradation process. Mitochondrial proteases, such as MPP and PARL, can cleave the PINK1 protein and prepare it for degradation (Thomas et al., 2014). When mitochondrial membrane potential gets changed; PINK1 protein senses it and stabilizes the damaged mitochondrial outer membrane. After that, PINK1 gets activated and recruits another protein Parkin from the cytosol. Parkin and PINK1 get phosphorylated for exerting their activities. Parkin then activates and interacts with various mitochondrial proteins with the help of Ubiquitin (Ub) (Sauvé et al., 2015). Mitochondrial adaptor proteins p97 and p62 then are ubiquitinated and recruited to facilitate clustering of mitochondria around perinuclear regions to selectively degrade the substrates via the proteasome system by a process referred to as autophagy or mitophagy (Lim et al., 2012). So, mutations in both the proteins and any one of the proteins can hamper the neuroprotective pathway at different stages through different molecular mechanisms and may abolish the pathway. PINK1 protein phosphorylates the Ser65 residue present in the Ubiquitin like (Ubl) domain of Parkin protein at its N-terminal end (Aguirre et al., 2017) (McWilliams et al., 2018). Interestingly, the activation of Parkin protein, its mitochondrial translocation and enzymatic functions are coupled together. PINK1 protein phosphorylates ubiquitin and the Ubl domain of Parkin exactly at the same positions that is the conserved Ser65 residue. Both of the phosphorylation processes are required for the activation of Parkin (Shiba-Fukushima et al., 2012) (Kazlauskaite et al., 2015) (Tang & Zhang, 2019). Parkin, an RBR (RING-in-between-RING (IBR)-RING) type E3 Ub ligase, has the ability to transfer the ubiquitin molecule from an E2 Ub-conjugating enzyme to the specific substrate (Dove & Klevit, 2017) (Metzger et al., 2014). The mechanism of the ubiquitin transfer is a two step process. Parkin’s RING1 domain first binds with the E2 enzyme to receive the Ub moiety from the E2 enzyme and forms a thioester intermediate by interacting with Parkin’s catalytic cystine (Cys431) residue (Iguchi et al., 2013). After that Ub is transferred from Parkin onto a lysine residue of a substrate protein (Durcan & Fon, 2015). So, PINK1 autophosphorylation and phosphorylation of ubiquitin and Ubl domain of Parkin are necessary for releasing the Ubl domain thereby activating the Parkin protein.

Mutation in the kinase domain of PINK1 protein can affect the autophosphorylation process and subsequently the activation of the Ubl domain of Parkin protein. The G309D mutation located in the kinase domain of PINK1 affects its interaction with the Ubl domain of Parkin (Iguchi et al., 2013) (Kumar et al., 2017). So, we considered this mutation for our current study. Along with this mutation we also chose the mutation P296L, as this mutation has highly pathogenic effect and is present in the Ins3 region of the kinase domain of PINK1 (Biswas & Bagchi, 2023).

In this work, we came up with a structural insight of the binding pattern of the PINK1-Parkin Ubl domain. For that, first both the wild type (WT) & mutated PINK1 proteins were docked with the Parkin Ubl domain. Then molecular dynamics simulations on the docked complexes were performed to analyze their cellular dynamics. Our simulations and *in silico* predictions would help to understand how the mutations could affect the Ubl domain binding and also could affect subsequent events. This PINK1-Parkin interaction is required for the activation of Parkin and its enzymatic functions. Importantly, our computational study will provide a clear view for a better understanding of the molecular mechanisms of PINK1-Parkin Ubl domain interaction.

## MATERIALS AND METHODS

### Structure Preparation

To conduct docking simulations, PINK1 and Parkin Ubl domain structures were retrieved from our previous work (Biswas et al., 2020) (Biswas & Bagchi, 2023). Molecular dynamics simulations of PINK1 kinase domain and Parkin Ubl domain were performed previously. We chose the average structure of the PINK1 from the stable zone of the simulation. In the similar way, Parkin Ubl domain structure was also retrieved. The Parkin Ubl domain is known to interact with PINK1 kinase domain’s Insertion 3 region. It is reported that a mutation in 309^th^ position can abolish the binding. In our previous work, we also identified some harmful mutations which could affect the activity of the kinase domain. From there, we selected the mutation at the 296^th^ position because it lies in the kinase domain’s Insertion 3 region. The mutants were built by using the ‘build mutant’ protocol of Discovery Studio 2.5 (DS2.5) platform.

### Model validation

SAVES 6.0 server (https://saves.mbi.ucla.edu/) server was used to evaluate the stereo-chemical properties of the structures. Ramachandran plots were drawn in Procheck tool. For the WT and mutant PINK1 kinase domain structures and Parkin Ubl domain, all the amino acid residues were present in the allowed regions of the Ramachandran plots (Ramachandran et al., 1963).

### Molecular Docking studies of PINK1 with Parkin Ubl domain

In this study, protein–protein docking simulation was performed to analyse the interactions between PINK1 kinase domain and Parkin Ubl domain. The WT PINK1 protein and two mutants of PINK1 protein were docked with Parkin Ubl domain in GRAMM-X software (Tovchigrechko & Vakser, 2006). GRAMM-X uses Fast Fourier Transform (FFT) to perform a rigid-body approach through applying smoothed Lennard–Jones potential, knowledge-based, and refinement stage scoring to discover the best surface match. From the docking studies, top ten docked poses were retrieved for further studies. The CHARMm force field was applied to the docked poses in DS 2.5 platform and optimized & re-ranked based on their binding free energy values. The docked pose having the lowest binding free energy was considered as the best docking pose. In our study, we specified both the receptor-ligand interacting residues which are as follows: Receptor (PINK1) binding site residues: Ser228, 357, Ser402 and Ligand (Parkin Ubl) binding site residues: Arg6, Asn8, Ile44, Lys48, Ser65, His68, Arg72 (Okatsuet al., 2018) (Rasool et al, 2018) (Rasool et al, 2022).

### Molecular Dynamics Simulation of the PINK1-Parkin Ubl domain docked complexes

Molecular dynamics simulations were performed for all three best docked complexes in GROMACS 5.1.5 package (Abraham et al., 2015). To generate the topology, CHARMm27 force field was applied to the docked complexes (system). In the next step, each of the systems was placed in a cubic periodic box maintaining adequate distance from the edges of the box. Then each of the systems was solvated and neutralized by adding SPC16 (simple point charge) water molecules and Na^+^ and Cl^-^ ions respectively. The simulations of each of the systems were performed in three steps,

1. energy minimization
2. heating,
3. equilibration and production run.

First of all, the built systems were energy minimized by the steepest descent method until the RMS gradient would reach 0.01 kcal/mole to relax the system. Then, using NPT (constant temperature and constant pressure ensemble) and NVT (constant temperature, constant volume) dynamics protocols the systems were heated and equilibrated under a constant pressure of 1 bar and a constant temperature of 300 K. After the equilibration procedure, the production run was performed for 180 ns.

### Trajectory Analysis

After completion of the molecular dynamics simulation MD trajectories were obtained. Using gmx_rmsd and gmx_rmsf tools the RMSD and RMSF of the three complexes were computed. The Rg and hydrogen bond analysis of the docked complexes were obtained using gmx_gyrate & gmx_hbond tools, respectively. The variations of the secondary structural elements of the protein complexes were analyzed using gmx_do_dssp tool. Graphs were plotted using Microsoft excel software and visualizations were done by using DS 2.5 server.

The principal component analysis (PCA) was done for all three docked complexes. The PCA was calculated by diagonalization and solution of the eigenvalues and eigenvectors for the covariance matrices obtained from the MD results. The eigenvectors and eigenvalues demonstrate the direction and the magnitude of the protein motion, respectively. Calculation of covariance matrix was performed using the tool gmx_covar. Further gmx_anaeig was applied for the calculation of overlap between the computed principal components and coordinates of the trajectory.

## RESULT AND DISCUSSION

### Structural analysis of the PINK1 and Parkin Ubl domain

To conduct the docking studies, the structures of the PINK1 kinase domain and Parkin Ubl domain were prepared. We only considered the PINK1 kinase domain and Parkin Ubl domain for this study because this interaction is very crucial for Parkin activity. So, from our previous simulation, both PINK1 and Parkin Ubl domain structures (Figure 1) were dumped from the stable zone of the MD simulations. Most favourable structures were selected, which also passed all the criteria for having a good model quality. These structures were then taken for docking studies. PINK1 mutants were built in D.S 2.5 platform and we also checked the Ramachandran plots.

**Figure 1.**
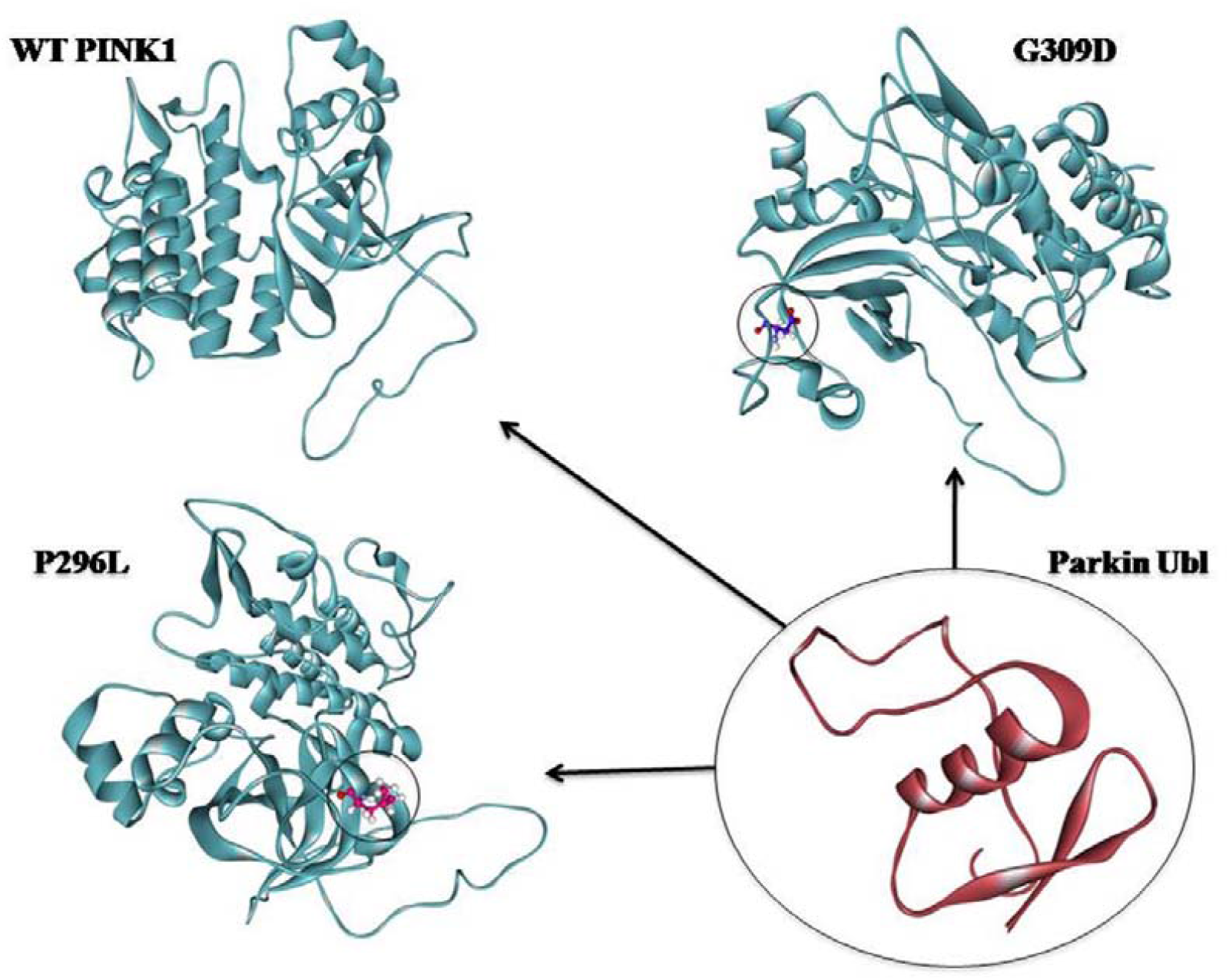
Three-dimensional structures of the PINK1 kinase domain, G309D & P296L mutants and Parkin Ubl domain are presented.

### Molecular Docking Simulations

The best ten docked pose for PINK1-Ubl complex according to the GRAMMX server were considered for further studies. In addition, according to the binding free energy score the docked complex with lowest binding free energy was considered as the best model (Table 1). The binding free energy of the WT PINK1-Ubl complex was -38.1635 kcal/mol. The lowest binding free energy of the P296L mutant-Ubl domain was -48.6201 kcal/mol and G309D mutant-Ubl domain was -26.2888 kcal/mol. The binding free energy shows that the interaction of G309D mutant with the Ubl domain of Parkin is less favourable with respect to the WT PINK1 and there were formations of less umber of hydrogen bonds. In contrast, P296L mutant had a higher binding free energy as compared to the WT PINK1 and due to this mutation unwanted bond formations would take place and this would create unfavourable situations.

**Table 1:**
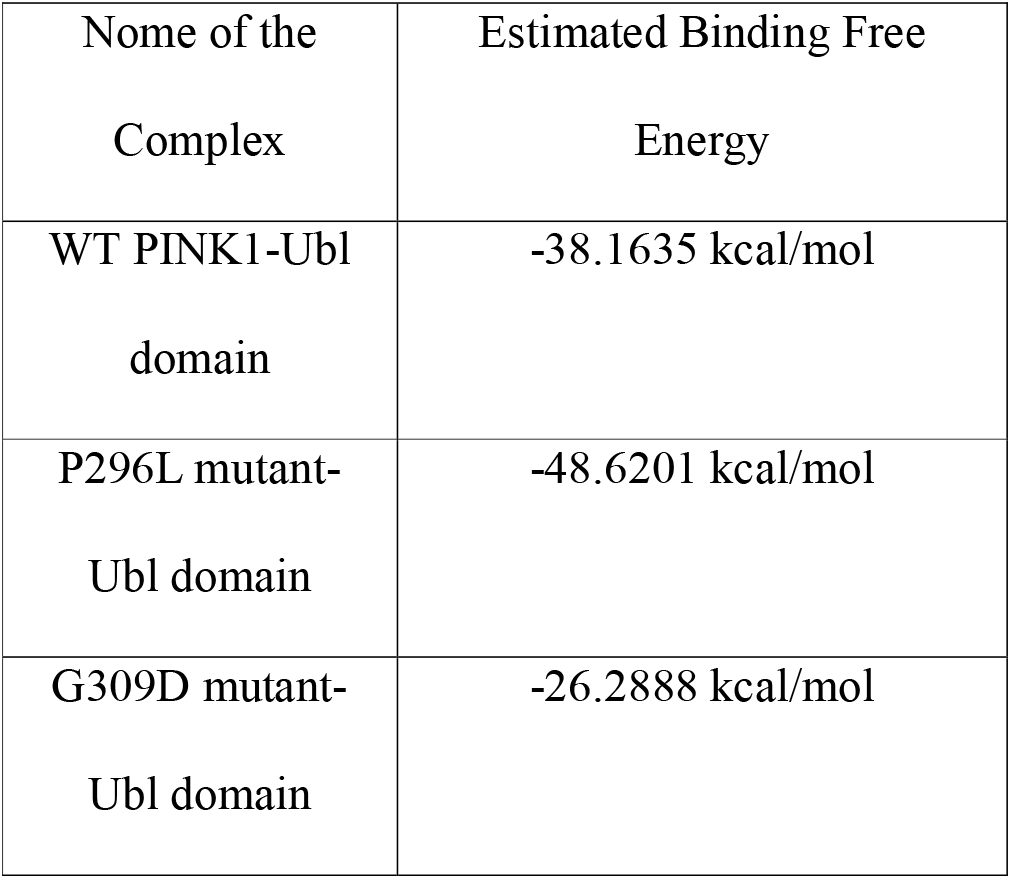
Best binding free energy of all the complex are presented in table 1.

### Molecular Dynamics Simulation

The root mean square deviation (RMSD) of the MD trajectory was analysed to check the stability of the protein complex over the MD simulation period. To analyse the dynamic stability of the docked complexes, RMSD profile of the backbone atoms of the amino acid residues were calculated and plotted against time (Figure 3). From the graph, it was observed that the WT docked complex achieved stability during the 180 ns course of MD simulation. For the G309D mutant complex the stability was fluctuating throughout the MD simulation and it did not achieve a stable conformation as compared to the WT docked complex. On the other hand, the mutant P296L would also show some fluctuations as compared to the WT docked complex.

**Figure 2.**
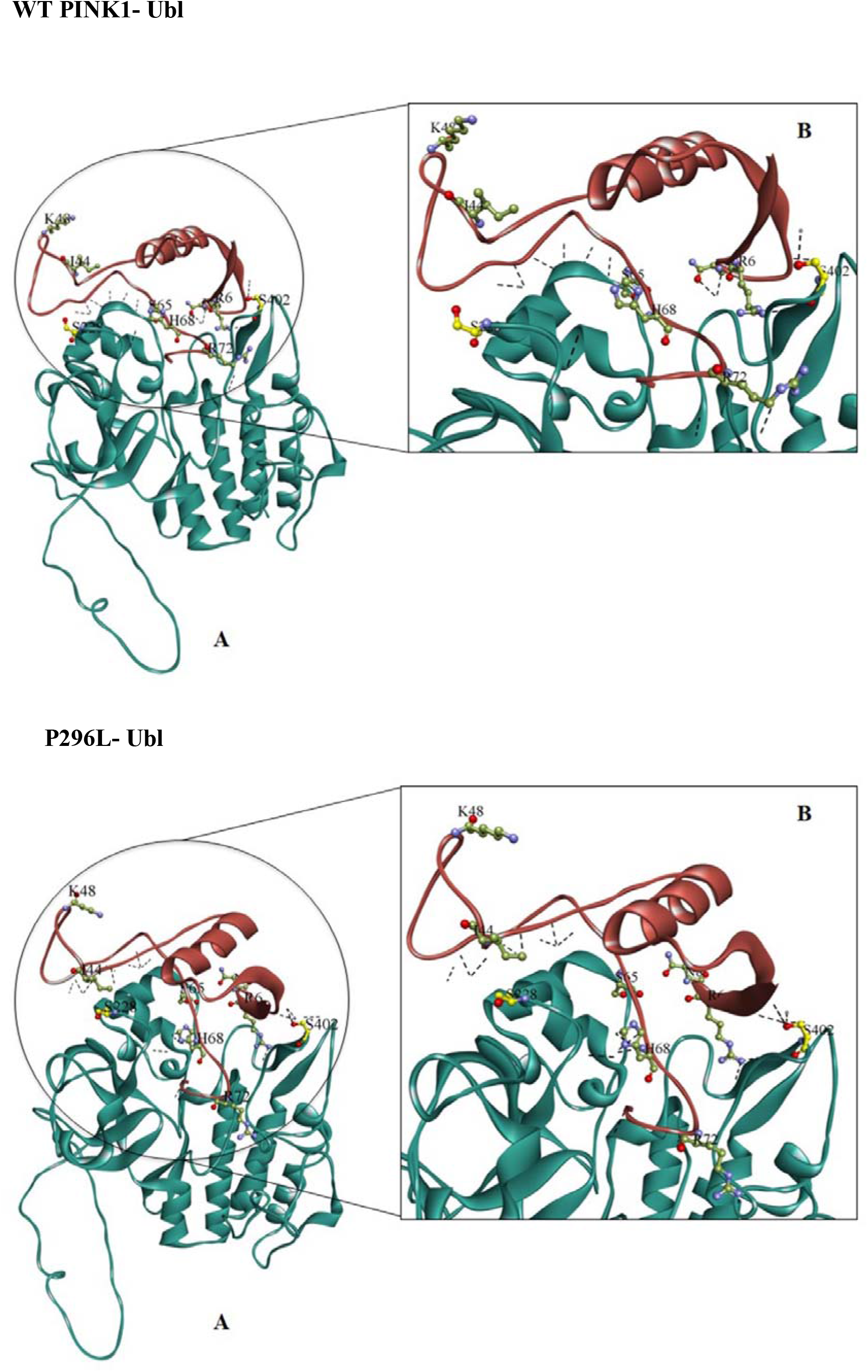
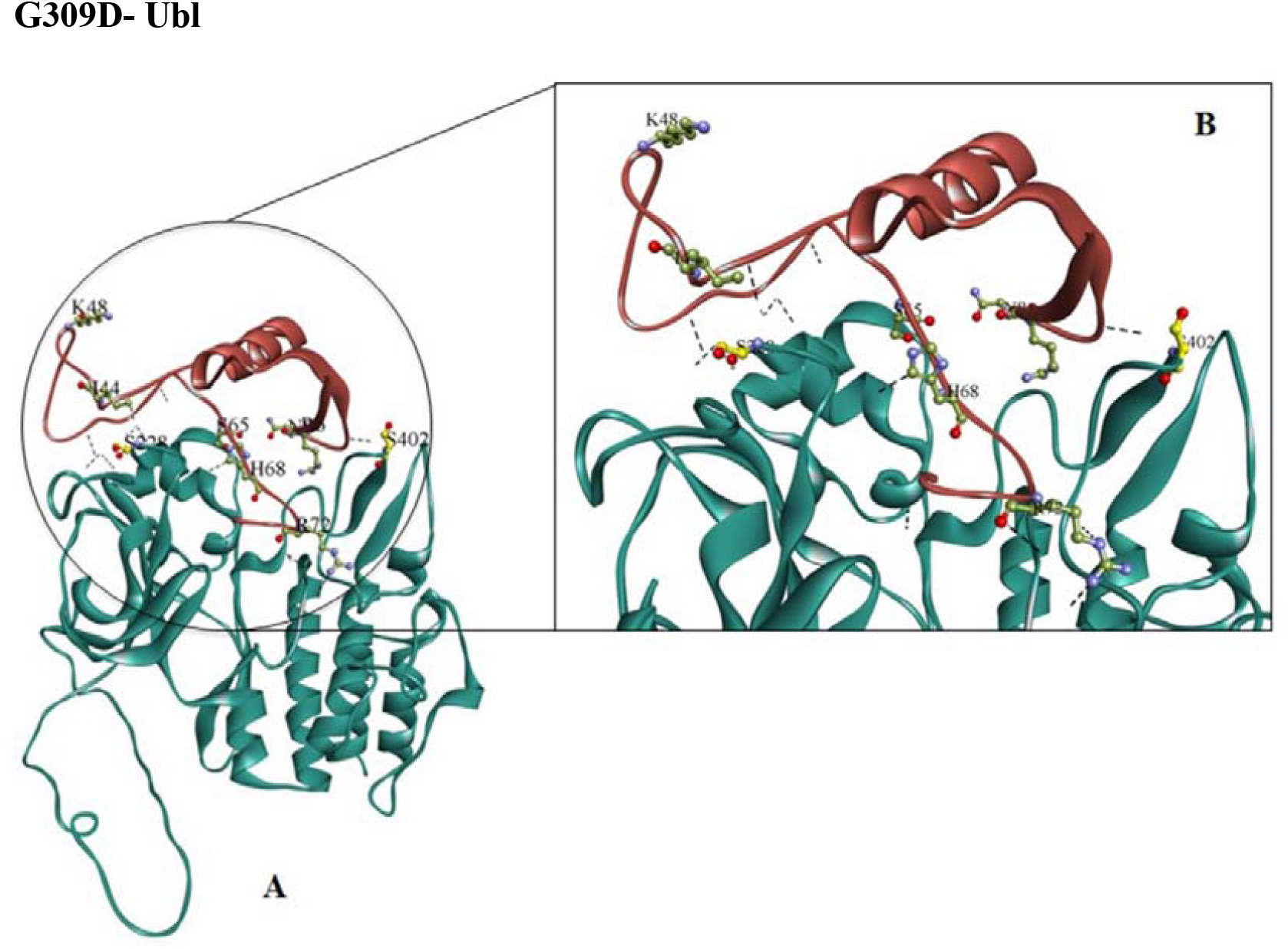
The three dimensional view of the docked complex and the main residues involved in the binding are visualised. In inset a detailed view of the hydrogen bonding interaction are presented.

**Figure 3.**
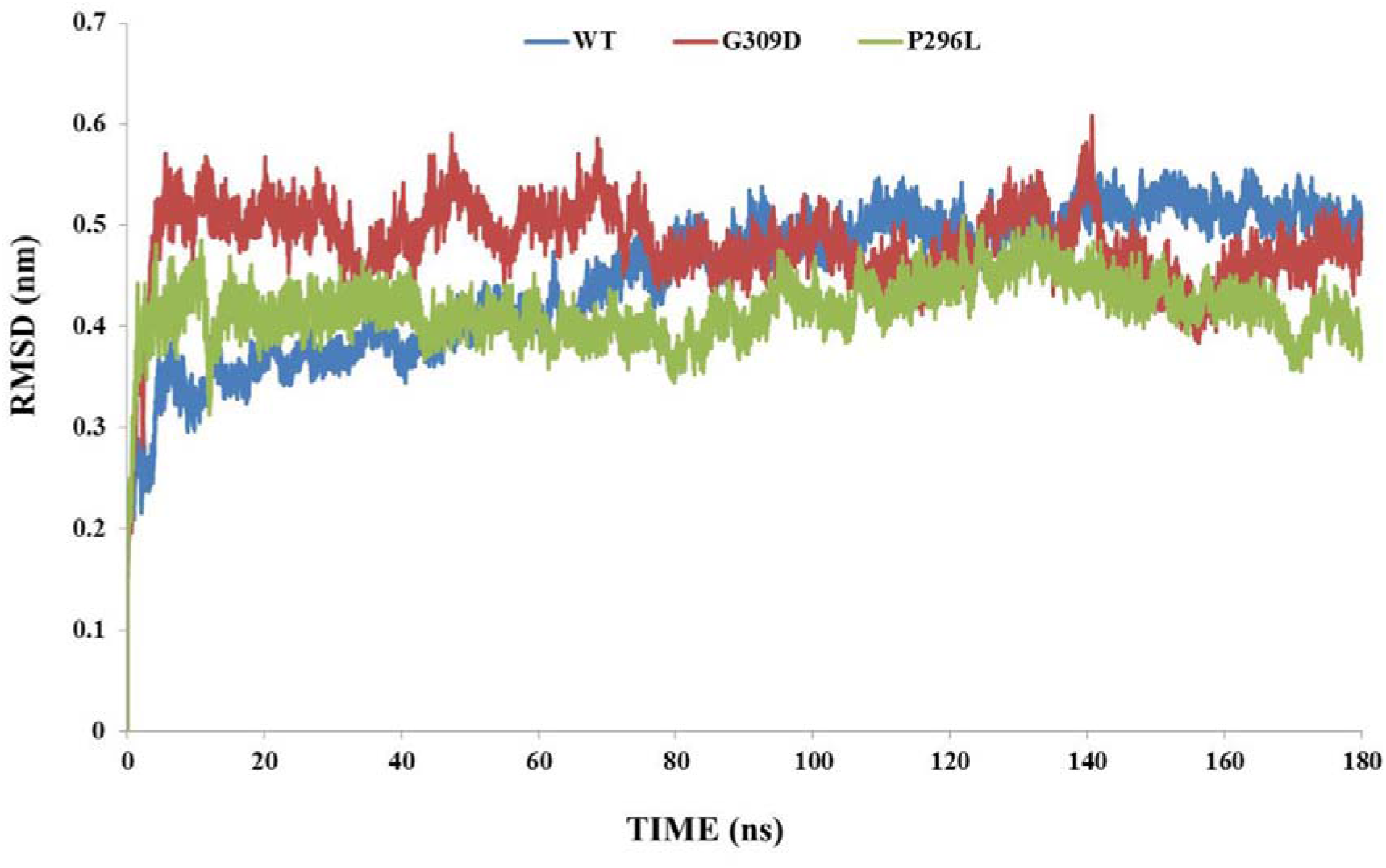
RMSD of the backbone atoms of the amino acid residues of the docked complexes during the simulation run of 180 ns. X-axis: time and Y-axis: RMSD (nm). Color scheme: WT (Blue), P296L (Green) and G309D (Red).

In order to observe the fluctuations of different amino acid residues, the RMSF plots of the C_α_ atoms were generated for all the docked complexes (Figure 4A & B). The highest fluctuations were observed in the region spanning 160-220 of the PINK1 protein. Insertion 3 is located in this region. So, due to the mutation this region is affected. The highest fluctuations were observed in the complex of P296L mutant with Ubl domain of Parkin among all the complexes. The level of fluctuations was less in G309D as compared to P296L. In contrast, Ubl domain’s residual fluctuations showed a reverse trend. The residual fluctuations are more throughout the course of the simulation in G309D mutation as compared to the WT and less in case of P296L mutation.

**Figure 4.**
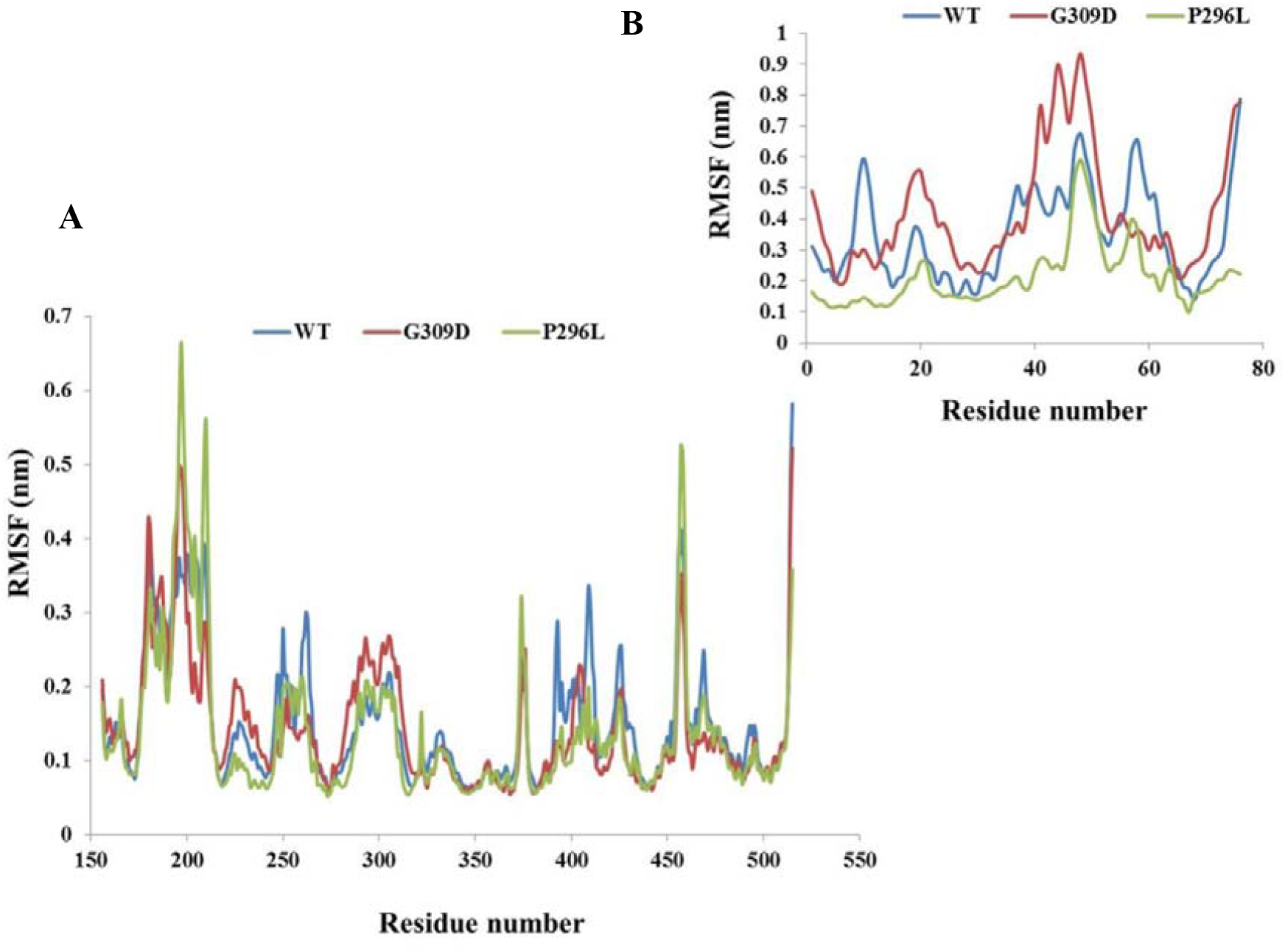
Root mean square fluctuation plots representing the fluctuations of the residues during the course of MD simulations for 180 ns of WTand mutant PINK1 kinase domains(A) and the Parkin Ubl domain (B); the X-axis: amino acid residue numbers and Y-axis: RMSF in nm. Color scheme: Native (Blue), P296L (Green) and G309D (Red).

The radius of gyration (Rg) was analysed to check the overall compactness of the docked complexes. If the Rg graph is stable over time-frame it can be predicted that the protein folding is stable. To check this, Rg vs. time graphs were plotted (Figure 5). The Rg value was stable in WT docked complex after some initial small fluctuations and maintains a stable compactness till the end of simulation time. The P296L mutant complex gains more stability which can hamper the proper protein activity. Moreover, in G309D, the complex failed to achieve a stable conformation throughout the simulation.

**Figure 5.**
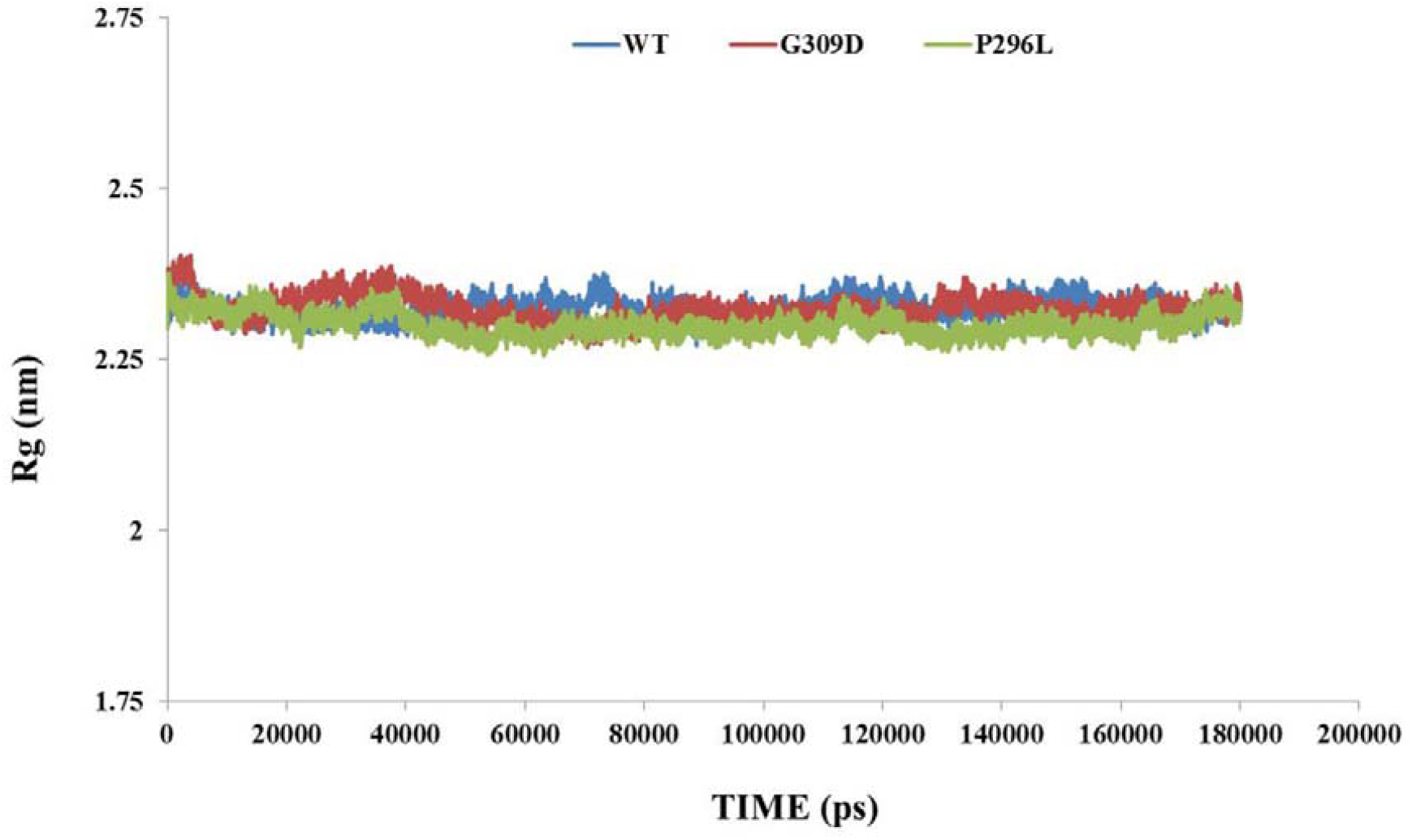
Radius of gyration (Rg) plot of WT and mutants over the 180 ns of MD simulation. X-axis: time in ps and Y-axis: Rg in nm. Color scheme: WT (Blue), P296L (Green) and G309D (Red).

The extent of hydrogen bonding in a protein complex depicts its stability. To achieve a stable binding between PINK1-Parkin Ubl domain the number of hydrogen bonds needs to remain stable throughout the MD simulation. In WT protein complex the number of hydrogen bonds was found to range between 6-27 approximately (Figure 6). In contrast, the extent of hydrogen bond formations was fluctuating initially in both the mutant complexes. The interacting residues which are participating in the hydrogen bonding interactions between PINK1 (A chain) and Parkin Ubl (C chain) are:

**Figure 6.**
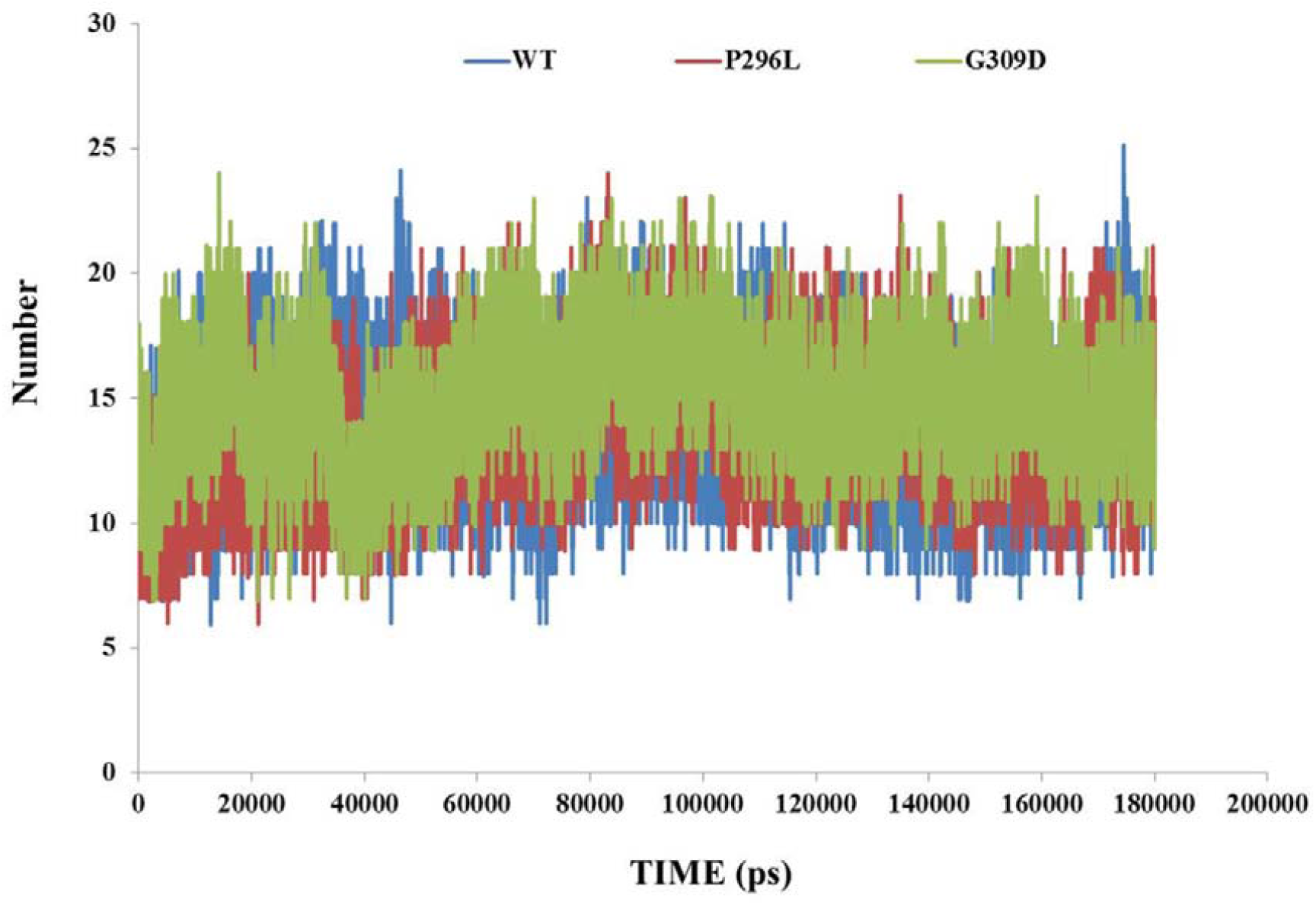
Plot of hydrogen bond analysis during the course of MD simulations. X-axis: number of the hydrogen bonds and Y-axis: Time in ps. Color scheme: Native (Blue), P296L (Red) and G309D (Green).

WT PINK1: Ser167, Lys206, Ile224, Ser229, Thr236, Tyr253, Arg259, Pro399, Ser401, Asp406 and Gly408

Ubl domain of Parkin: Arg6, Gly12, Arg42, Thr55, Gln57, Cys59, Asp62, Ile66, Gln71, Trp74 and Arg75.

The residues and the bonds are changed in the docked complexes of the mutants. In G309D mutant complex the hydrogen bond formation is less, whereas in P296L mutant complex more hydrogen bond formation occurs with respect to the WT docked complex (Figure 10).

The hydrogen bonds were visualized using the tool LigPlus and depicted in Figure 10.

To elucidate the dynamical properties of the docked complexes the fluctuations and structural motions of the atoms of the amino acid residues in the complexes were measured by PCA or Essential dynamics (ED) analysis. The atomic motions were found to be largely different for all the complexes. For, the WT complex a positive variation mainly along the PC1 axis was noted. On the other hand, for the mutant P296L, the trend is almost reversed. In this case, the dominant motions were found to be along the negative direction of the PC2. The extents of the atomic movements were almost completely opposite for the mutant G309D. All these would signify that the mutations in PINK1 would render changes in the complexes to a large extent. A larger conformational change was observed in the G309D mutation compared to the WT.

Colour code: Red: High positive correlation; Blue: Strong negative correlation

The dynamic cross-correlation maps (Figure 8) of the different protein complexes would reveal that atomic movements of the components of the WT complex are somewhat well-correlated as indicated by the red lines. On the other hand, the correlation between the atomic movements was much less for the mutant G309D. In other words, among all the cases, the mutation G309D would create a different structural environment in such a way that the mode of interactions between the protein components in the complexes would change from the WT to a large extent.

**Figure 7.**
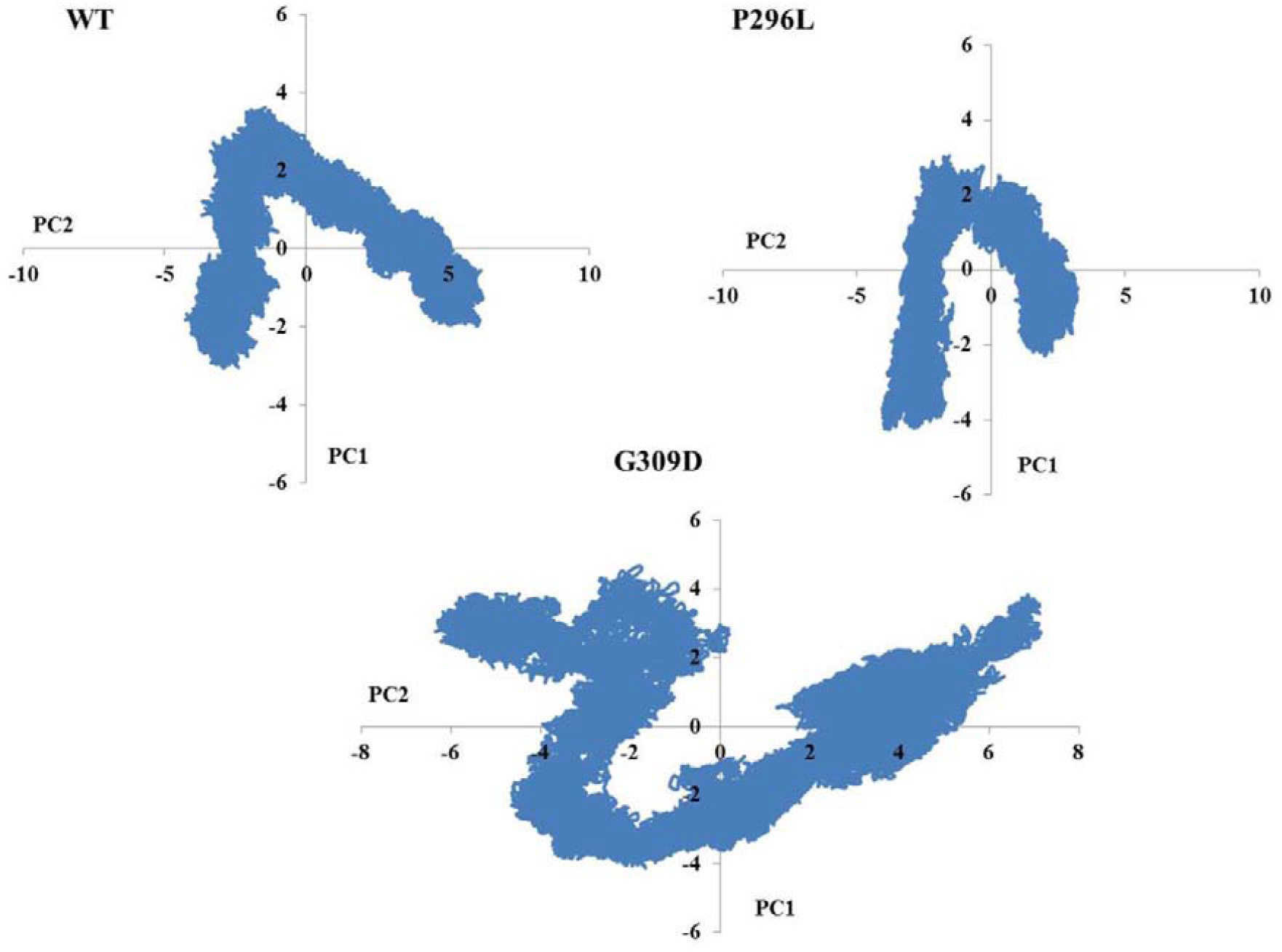
PCA analysis of the WT and mutant docked complexes during the course of MD simulations.

**Figure 8.**
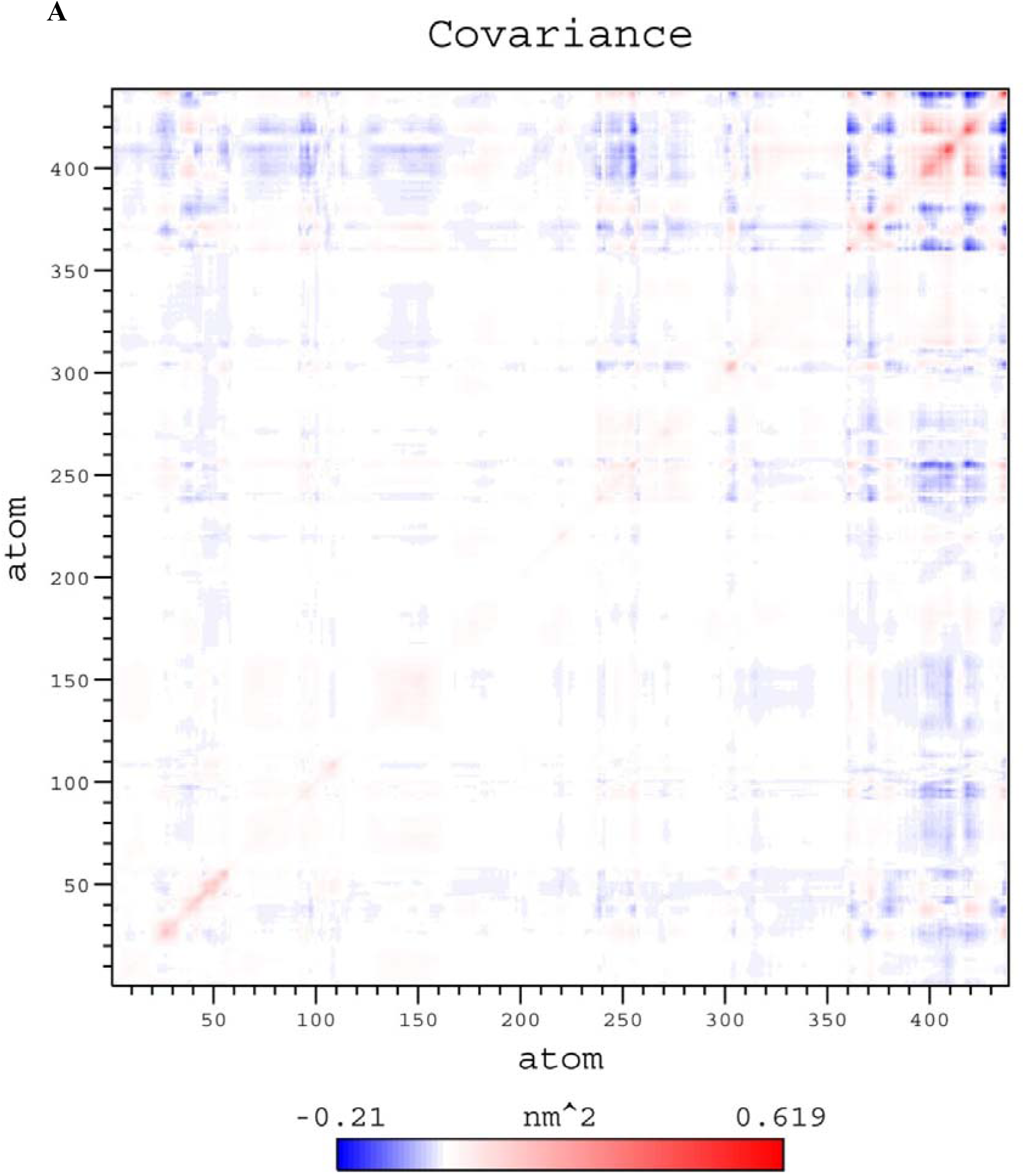
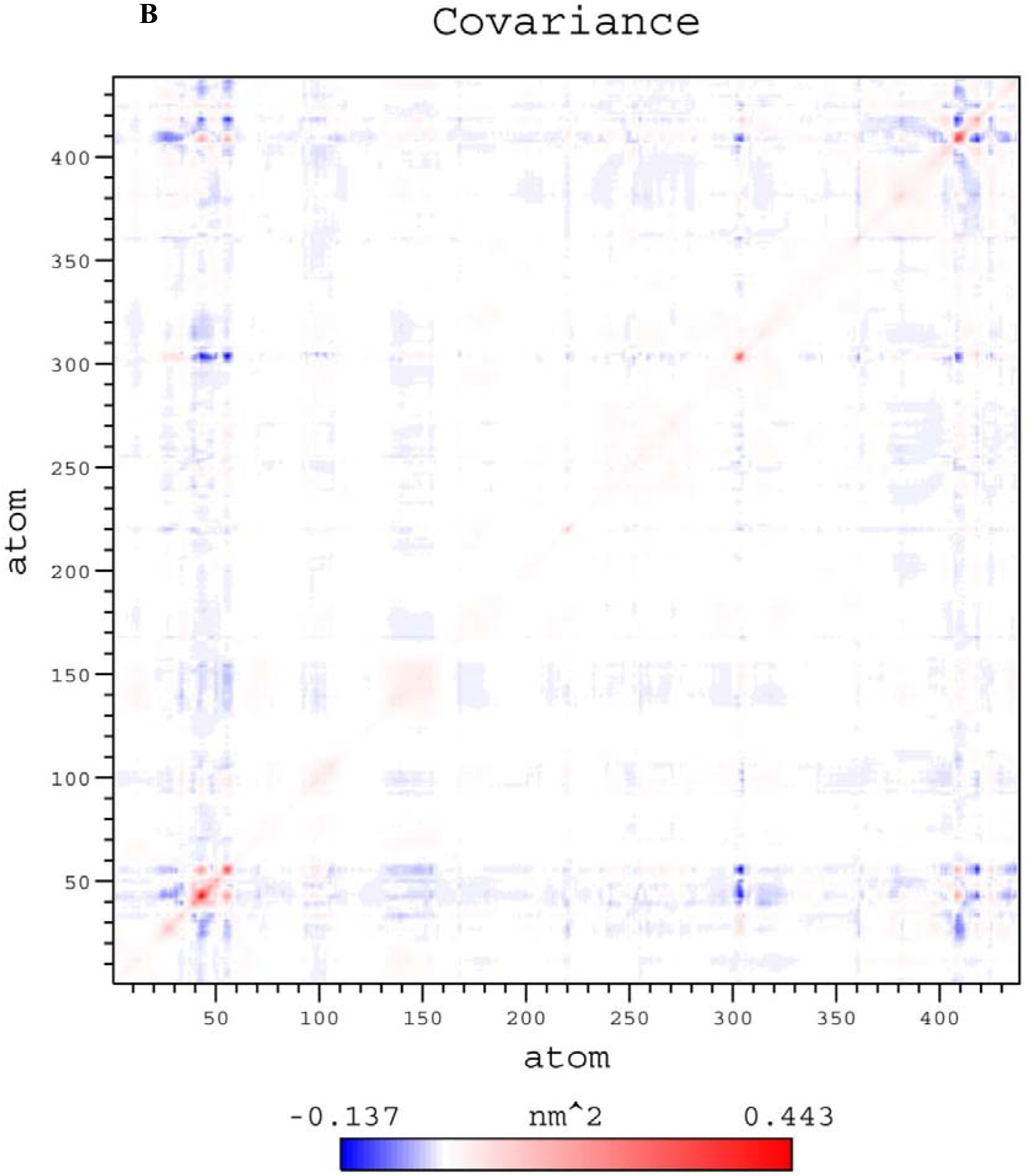
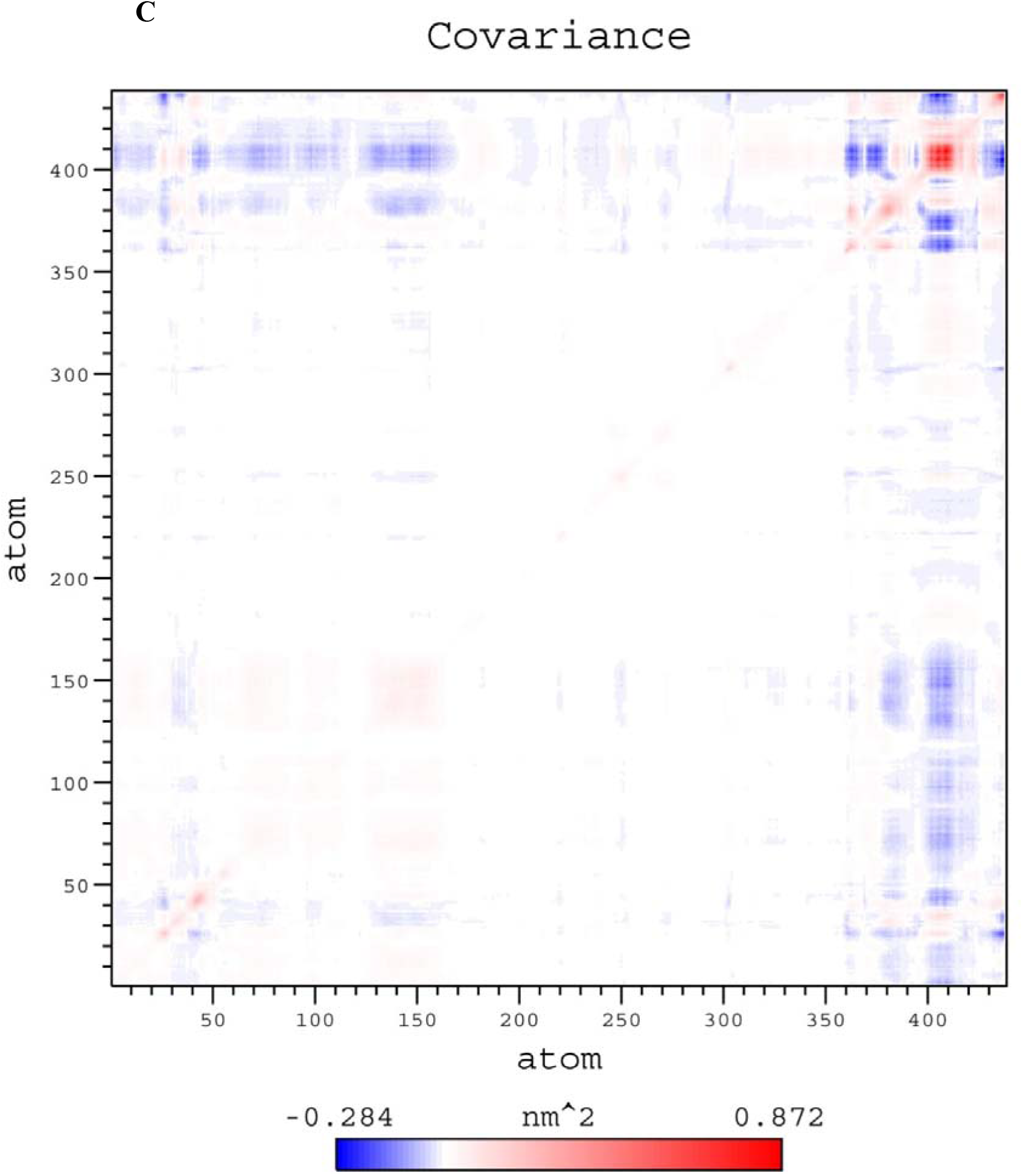
The dynamic cross-correlation map (DCCM) of the WT PINK1-Ubl complex (A), mutant PINK1 (P296L)-Ubl complex (B) and mutant PINK1 (G309D)-Ubl complex (C).

Secondary structure analysis was done by DSSP tool to check the secondary structural changes throughout the simulations (Figure 9). The fluctuations of the secondary structural elements in the G309D complex were found to be more than the other two complexes.

**Figure 9.**
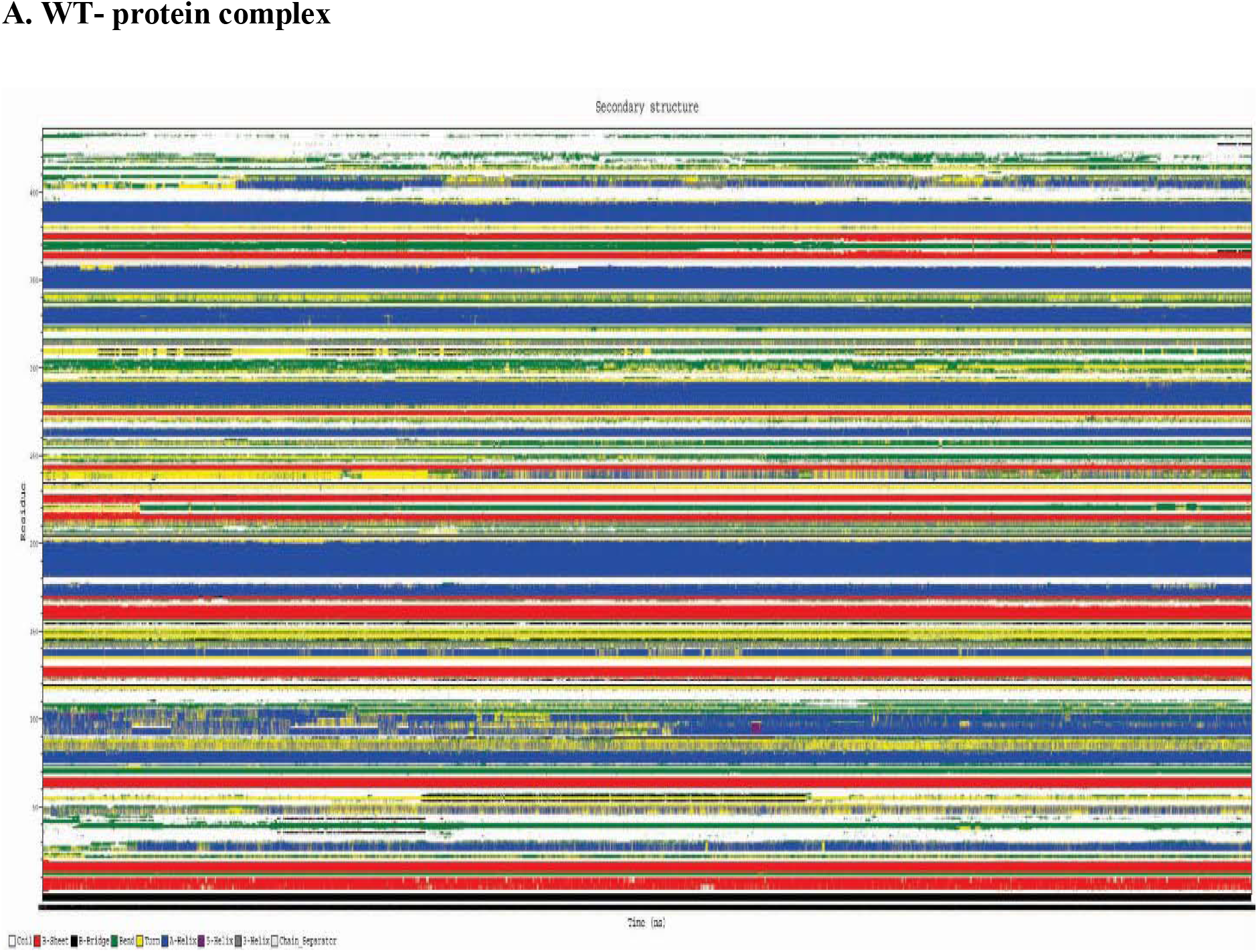
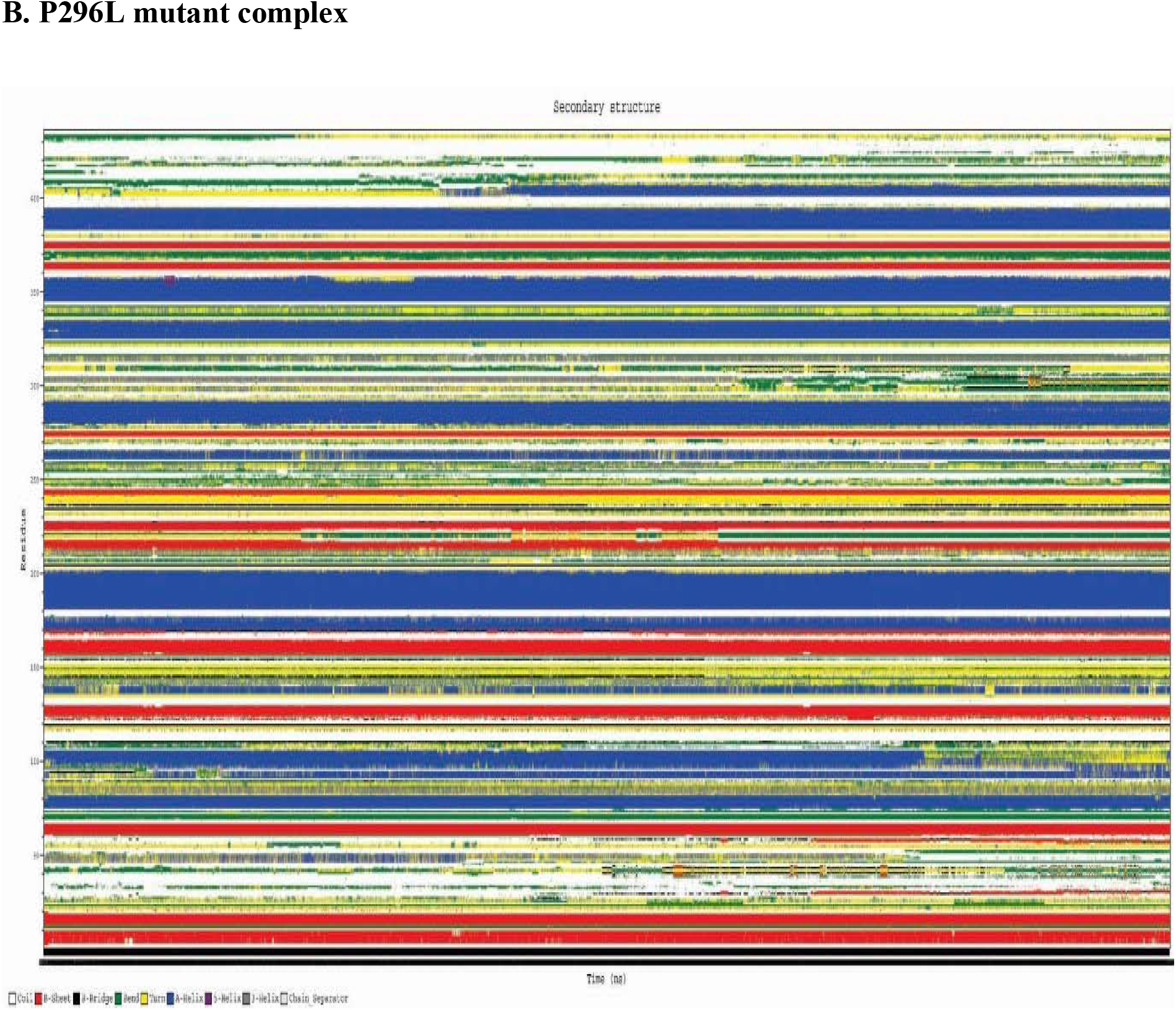
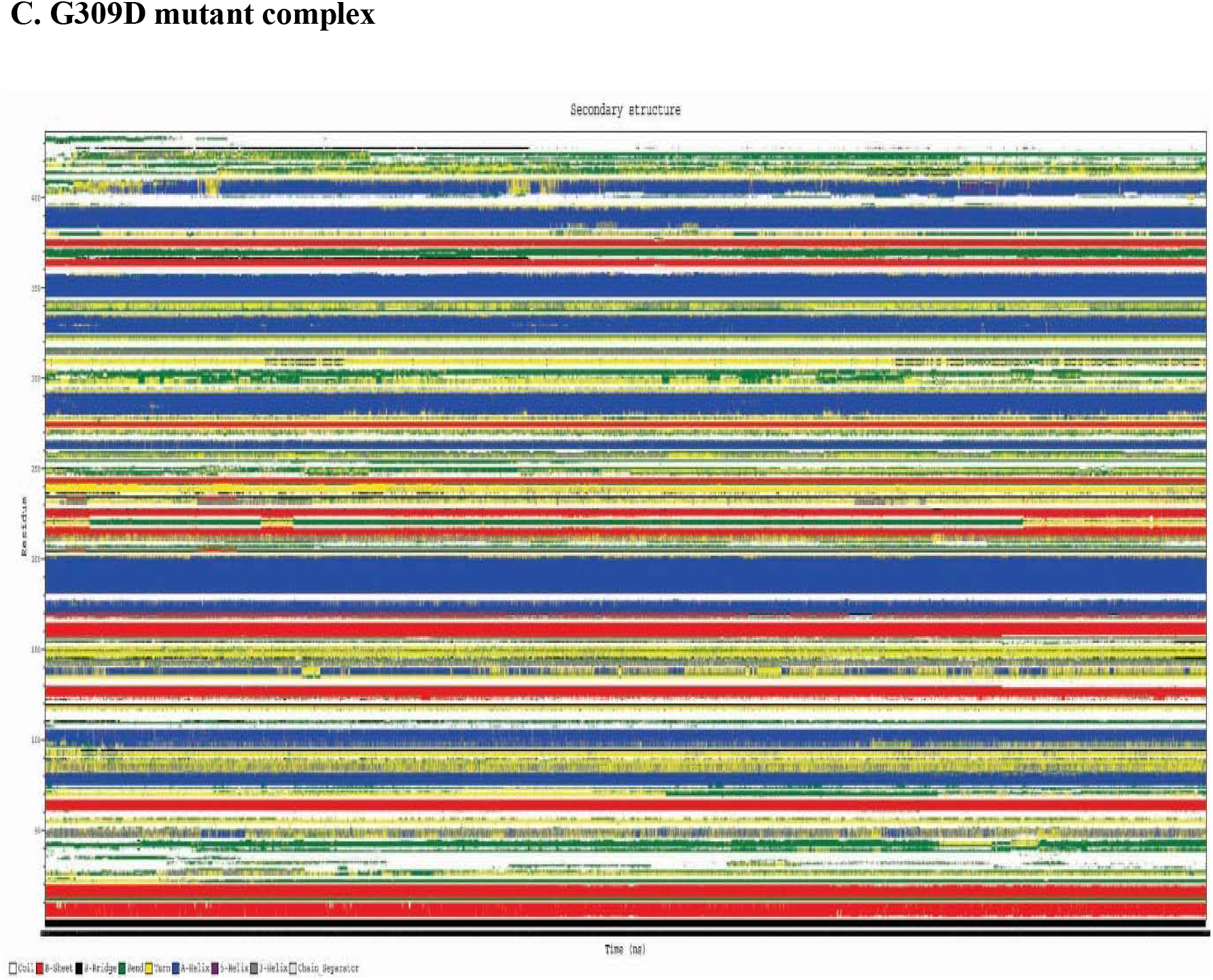
Changes in the secondary structural elements of PINK1-Ubl complexes. A: WT complex B: P296L mutant complex C: G309D mutant complex

**Figure 10.**
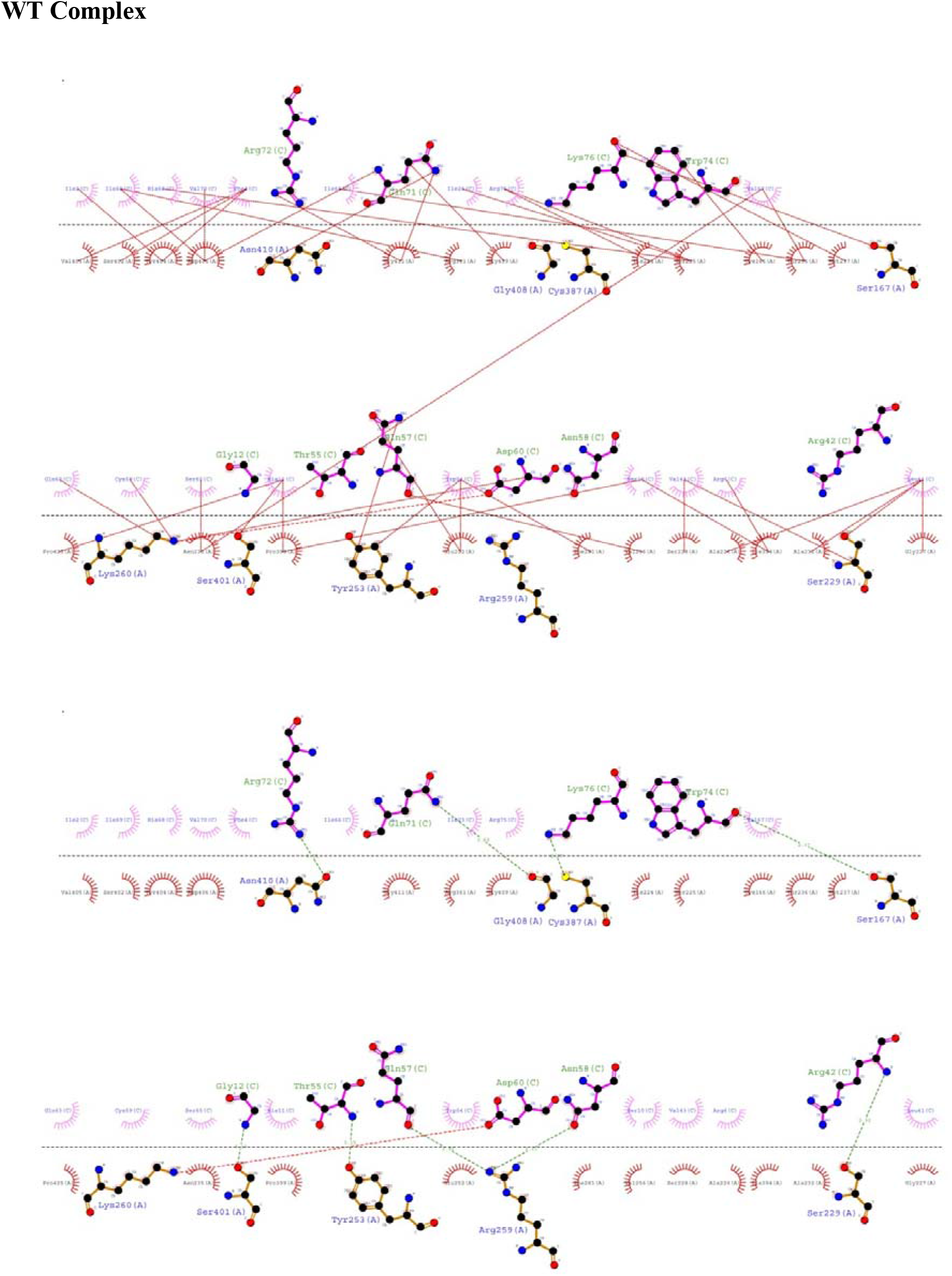
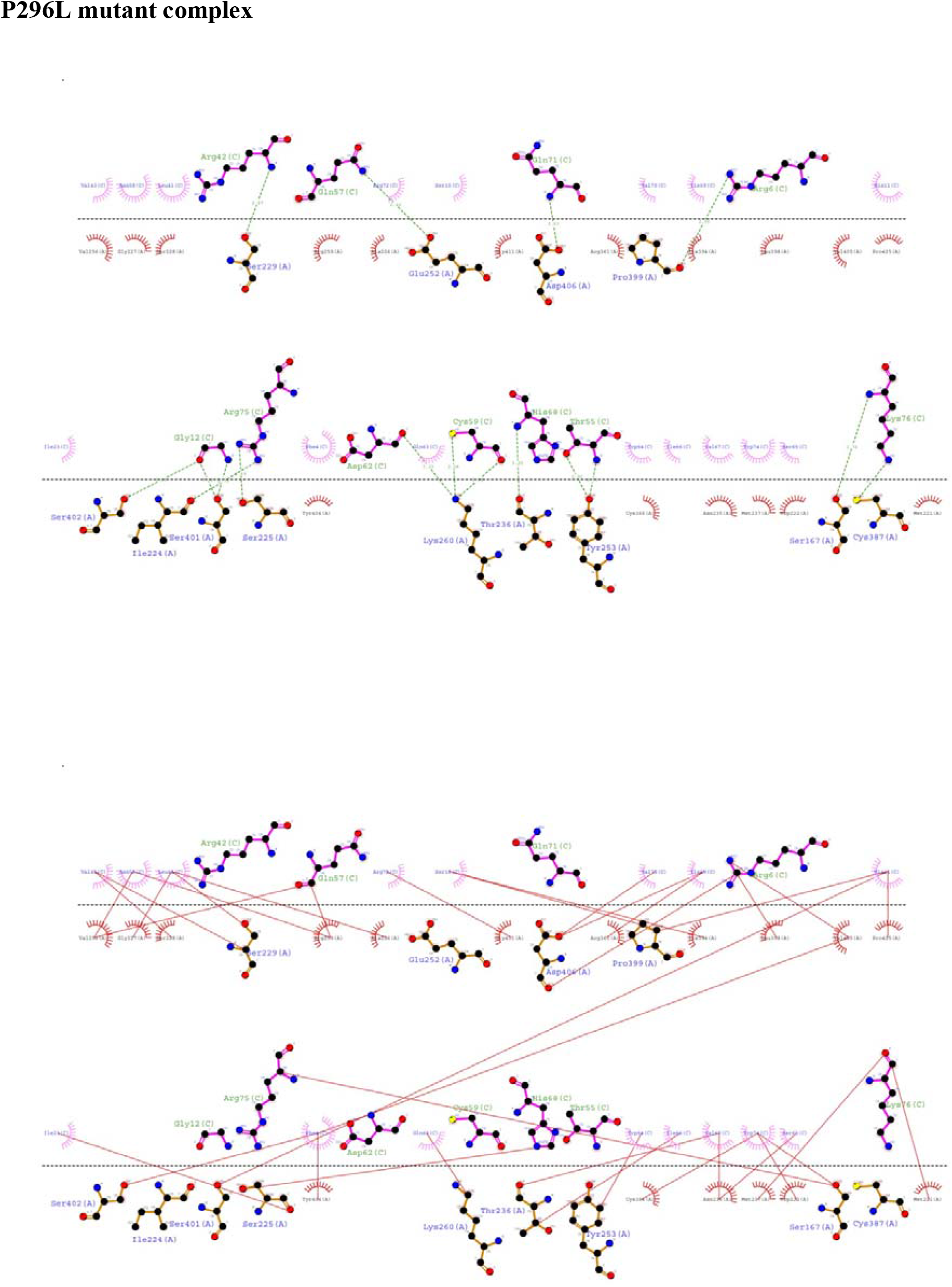
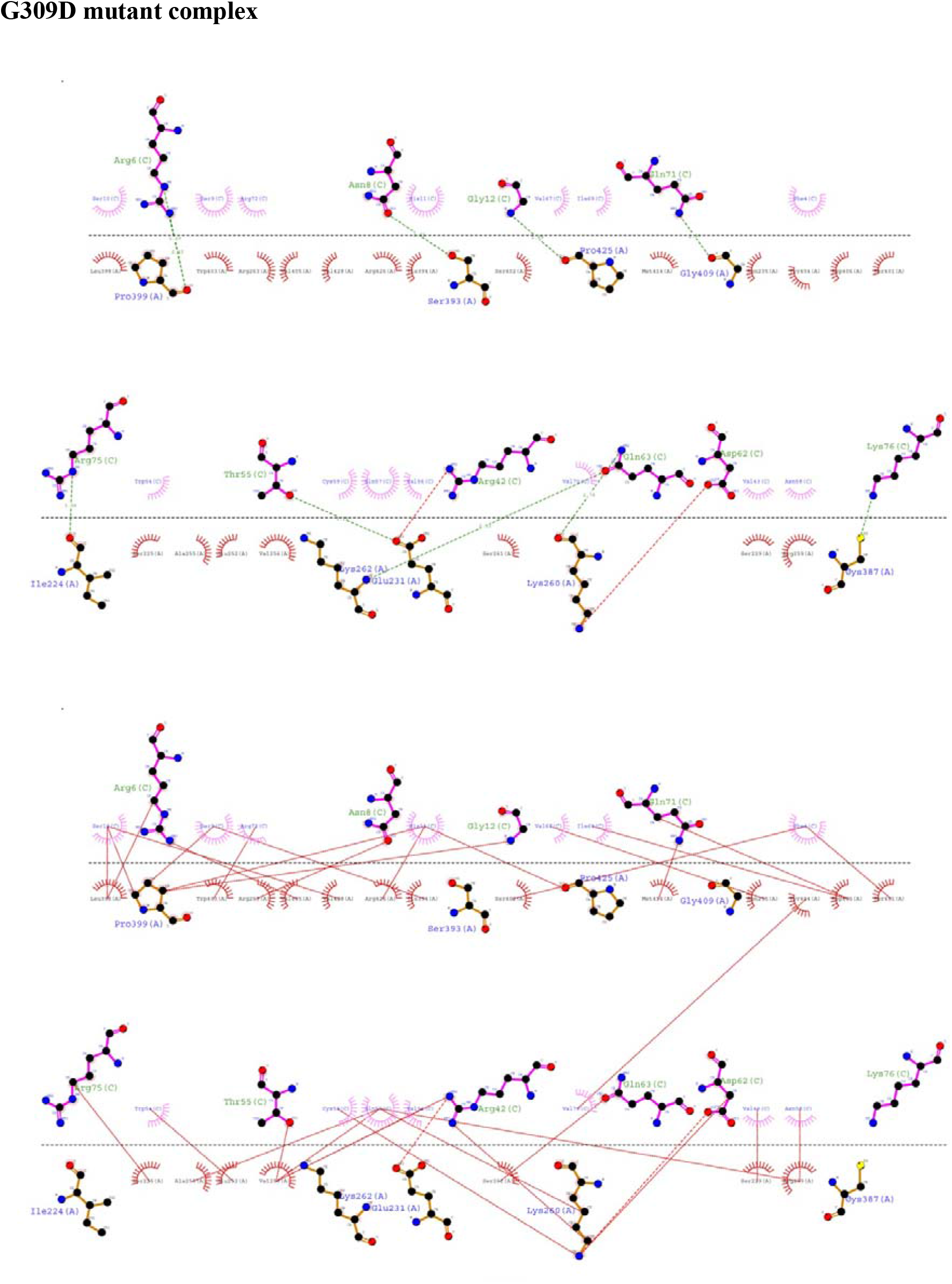
Dimplot analysis results. Hydrogen and hydrophobic interactions are visualised in dimplot for a better understanding of the mode of non-covalent interactions in the docked complexes.

## DISCUSSION

We first identified the mutations of the insertion3 region in PINK1 kinase domain which will affect the PINK1 and Parkin Ubl domain interactions necessary for the activation of Parkin. In addition, this domain interaction is required for activation of its enzymatic function. Therefore, to study the mutational effect on this interaction we conducted docking studies. All the docked complexes were energy minimized in D.S 2.5 and the binding free energy values were calculated. Based on lowest binding free energy the best docked complex was selected in each case.

The protein–protein complex having minimum binding free energy was further subjected to MD simulations analysis till 180 ns. The native complex achieved its stability at 180ns but the mutants are not stable at this point. On the other hand, from the H-bonds analysis, we were able to determine the stable pattern of interactions between PINK1 and Parkin Ubl during course of the simulation.

## CONCLUSION

This study will give structural insights into the PINK1-Parkin interactions. Molecular docking and molecular dynamics simulation studies were conducted to identify structural insight into the interaction pattern between PINK1 and Parkin Ubl domain. Our study showed how the mutations in the PINK1 protein’s insertion3 region would affect its interaction with Parkin Ubl domain. Interestingly, due to the G309D mutation, the PINK1-Parkin Ubl domain interaction was hampered and it also reflected in the binding free energy. This work provides the binding interactions between the WT PINK1 & mutated PINK1 with Parkin Ubl domain will be useful for further related studies, including understanding the molecular mechanisms of the Parkin Ubl domain phosphorylation and Parkin protein activation. This study will help to understand the molecular mechanism behind the PINK1 mediated Parkin activation.

## ACKNOWLEDGEMENT

The authors acknowledge the financial support from the DBT funded Bioinformatics Infrastructure Facility Center (Sanction no.: BT/PR40162/BTIS/137/48/2022 sanctioned to Prof. Angshuman Bagchi). Sima Biswas receives funding from University of Kalyani. Other funding sources are DST PURSE, UGC-SAP.

## REFERENCES

Abraham, M. J., Murtola, T., Schulz, R., Páll, S., Smith, J. C., Hess, B., & Lindahl, E. (2015). GROMACS: High performance molecular simulations through multi-level parallelism from laptops to supercomputers. SoftwareX, 1, 19–25.

Agarwal, S., & Muqit, M. M. (2022). PTEN-induced kinase 1 (PINK1) and Parkin: unlocking a mitochondrial quality control pathway linked to Parkinson’s disease. Current opinion in neurobiology, 72, 111–119.

Aguirre, J. D., Dunkerley, K. M., Mercier, P., & Shaw, G. S. (2017). Structure of phosphorylated UBL domain and insights into PINK1-orchestrated parkin activation. Proceedings of the National Academy of Sciences, 114(2), 298–303.

Biswas, S., & Bagchi, A. (2023). Analysis of the structural dynamics of the mutations in the kinase domain of PINK1 protein associated with Parkinson’s disease. Gene, 147183.

Biswas, S., Roy, R., Biswas, R., & Bagchi, A. (2020). Structural analysis of the effects of mutations in Ubl domain of Parkin leading to Parkinson’s disease. Gene, 726, 144186.

Cookson, M. R. (2012). Parkinsonism due to mutations in PINK1, parkin, and DJ-1 and oxidative stress and mitochondrial pathways. Cold Spring Harbor perspectives in medicine, 2(9), a009415.

Deas, E., Wood, N. W., & Plun-Favreau, H. (2011). Mitophagy and Parkinson’s disease: the PINK1–parkin link. Biochimica et BiophysicaActa (BBA)-Molecular Cell Research, 1813(4), 623–633.

Dove, K. K., & Klevit, R. E. (2017). RING-between-RING E3 ligases: emerging themes amid the variations. Journal of molecular biology, 429(22), 3363–3375.

Durcan, T. M., & Fon, E. A. (2015). The three ‘P’s of mitophagy: PARKIN, PINK1, and post–translational modifications. Genes & development, 29(10), 989–999.

Ge, P., Dawson, V. L., & Dawson, T. M. (2020). PINK1 and Parkin mitochondrial quality control: A source of regional vulnerability in Parkinson’s disease. Molecular neurodegeneration, 15, 1–18.

Iguchi, M., Kujuro, Y., Okatsu, K., Koyano, F., Kosako, H., Kimura, M., Suzuki, N., Uchiyama, S., Tanaka, K. & Matsuda, N. (2013). Parkin-catalyzed ubiquitin-ester transfer is triggered by PINK1-dependent phosphorylation. Journal of biological chemistry, 288(30), 22019–22032.

Iguchi, M., Kujuro, Y., Okatsu, K., Koyano, F., Kosako, H., Kimura, M., Suzuki, N., Uchiyama, S., Tanaka, K & Matsuda, N. (2013). Parkin-catalyzed ubiquitin-ester transfer is triggered by PINK1-dependent phosphorylation. Journal of biological chemistry, 288(30), 22019–22032.

Kazlauskaite, A., Martínez-Torres, R. J., Wilkie, S., Kumar, A., Peltier, J., Gonzalez, A., Johnson, C., Zhang, J., Hope, A.G., Peggie, M., Trost, M., & Muqit, M. M. (2015). Binding to serine 65-phosphorylated ubiquitin primes Parkin for optimal PINK 1-dependent phosphorylation and activation. EMBO reports, 16(8), 939–954.

Kitada, T., Asakawa, S., Hattori, N., Matsumine, H., Yamamura, Y., Minoshima, S., Yokochi, M., Mizuno, Y. & Shimizu, N. (1998). Mutations in the parkin gene cause autosomal recessive juvenile parkinsonism. nature, 392(6676), 605–608.

Kumar, A., Tamjar, J., Waddell, A. D., Woodroof, H. I., Raimi, O. G., Shaw, A. M., Peggie, M., Muqit, M.M, & van Aalten, D. M. (2017). Structure of PINK1 and mechanisms of Parkinson’s disease-associated mutations. Elife, 6, e29985.

Lim, K. L., Ng, X. H., Grace, L. G. Y., & Yao, T. P. (2012). Mitochondrial dynamics and Parkinson’s disease: focus on parkin. Antioxidants & redox signaling, 16(9), 935–949.

Lüthy, R., Bowie, J. U., & Eisenberg, D. (1992). Assessment of protein models with three-d Ramachandran, G.N., Ramakrishnan, C. and Sasisekharan, V. (1963) Stereochemistry of Polypeptide Chain Configurations. Journal of Molecular Biology, 7, 95-99.imensional profiles. Nature, 356(6364), 83–85.

McWilliams, T. G., Barini, E., Pohjolan-Pirhonen, R., Brooks, S. P., Singh, F., Burel, S., Balk, K., Kumar, A., Montava-Garriga, L., Prescott, A.R. and Hassoun, S.M. & Muqit, M. M. (2018). Phosphorylation of Parkin at serine 65 is essential for its activation in vivo. Royal Society Open Biology, 8(11), 180108.

Metzger, M. B., Pruneda, J. N., Klevit, R. E., & Weissman, A. M. (2014). RING-type E3 ligases: master manipulators of E2 ubiquitin-conjugating enzymes and ubiquitination. Biochimica et BiophysicaActa (BBA)-Molecular Cell Research, 1843(1), 47–60.

Okatsu, K., Sato, Y., Yamano, K., Matsuda, N., Negishi, L., Takahashi, A., Yamagata, A., Goto-Ito, S., Mishima, M., Ito, Y., Oka, T., & Fukai, S. (2018). Structural insights into ubiquitin phosphorylation by PINK1. Scientific reports, 8(1), 10382.

Quinn, P. M., Moreira, P. I., Ambrósio, A. F., & Alves, C. H. (2020). PINK1/PARKIN signalling in neurodegeneration and neuroinflammation. Actaneuropathologica communications, 8(1), 1–20.

Rasool, S., Soya, N., Truong, L., Croteau, N., Lukacs, G. L., & Trempe, J. F. (2018). PINK 1 autophosphorylation is required for ubiquitin recognition. EMBO reports, 19(4), e44981.

Sauvé, V., Lilov, A., Seirafi, M., Vranas, M., Rasool, S., Kozlov, G., Sprules, T., Wang, J., Trempe, J.F. & Gehring, K. (2015). A Ubl/ubiquitin switch in the activation of Parkin. The EMBO journal, 34(20), 2492–2505.

Shiba-Fukushima, K., Imai, Y., Yoshida, S., Ishihama, Y., Kanao, T., Sato, S., & Hattori, N. (2012). PINK1-mediated phosphorylation of the Parkin ubiquitin-like domain primes mitochondrial translocation of Parkin and regulates mitophagy. Scientific reports, 2(1), 1–8.

Tang, C., & Zhang, W. P. (2019). How phosphorylation by PINK1 remodels the ubiquitin system: A perspective from structure and dynamics. Biochemistry, 59(1), 26–33.

Thomas, R. E., Andrews, L. A., Burman, J. L., Lin, W. Y., & Pallanck, L. J. (2014). PINK1-Parkin pathway activity is regulated by degradation of PINK1 in the mitochondrial matrix. PLoS genetics, 10(5), e1004279.

Tovchigrechko, A., & Vakser, I. A. (2006). GRAMM-X public web server for protein–protein docking. Nucleic acids research, 34(Suppl_2), W310–W314.

